# Radiocarbon dating of bird and mammal eye lenses reveals long-term chronology

**DOI:** 10.64898/2026.09.15.751752

**Authors:** Emi Hasegawa, Jun Matsubayashi, Kazuki Miura, Yumi Yamanashi, Sinta Maharani, Ichiro Tayasu

## Abstract

Time-series isotope analysis using eye lenses has recently gained attention as a low-cost method for reconstructing long-term information from a single sampling event, with potential applications across diverse taxonomic groups. However, information from eye lenses in birds and mammals remains limited, particularly during early growth stages, and the eye-lens time axis over extended periods should be reconstructed. This study measured the radiocarbon (^14^C) isotope ratios of eye lenses to examine the temporal framework recorded in the eye lenses of birds and mammals. The calibrated ^14^C ages of the central lens sections closely matched the known birth years or years preceding their introduction to Japan. ^14^C ages increased progressively from the center toward the periphery of the eye lens. The Siberian tiger allowed information reconstruction spanning from 2002.4 ± 1.6 to 2017.8 ± 2.0, whereas the ring-tailed lemur yielded reconstructed ages from 2000.2 ± 1.6 to 2019.2 ± 0.0. Calendar ages for all individuals after approximately 2019 could not be resolved using the radiocarbon calibration approach applied in this study. The sulphur-crested cockatoo demonstrated the longest reconstructed time span, covering 27.4 years from 1968.7 ± 1.3 to 1996.1 ± 1.4. Our results indicate that eye lenses can reconstruct long-term information spanning much of the lifespan in mammals examined, whereas avian lenses can preserve information spanning several years to decades after birth. Lens growth patterns may differ according to life history; thus, future studies should examine a wider range of species and taxonomic groups with diverse life histories.

## 1 Introduction

Time-series isotope analysis using eye lenses has recently gained attention because it is a low-cost method that collects information to replace long-term monitoring from the past with just a single sampling (Matsubayashi et al. 2025). The eye lens has a structure in which new lens fiber cells are continuously formed in a concentric, annual ring-like pattern over older cells. As the cells mature, they lose organelles, including the nucleus and mitochondria, which are necessary for proliferation and metabolism (Bassnett 2002). New material accretes from the center (nucleus) toward the outer cortex; thus, radial subsampling of the lens can provide a chronological sequence suitable for reconstructing ontogenetic shifts in habitat use and diet. The retrospective isotope analysis of the eye lens has been applied largely to questions in early life history and natal origin. Notable applications include the radiocarbon analysis of eye lens cores to demonstrate exceptional longevity in Greenland sharks (Nielsen et al. 2016) and the use of eye lens isotope ratios as natural markers to distinguish hatchery stocks from naturally reproduced trout (Rosinski et al. 2023).

The original eye lens isotope approach was developed using fresh lenses (Wallace et al. 2014). However, back-calculation is challenging when a given lens section was deposited because the peripheral layers could not be subdivided at sufficiently high resolution. Further, the method was less readily transferable beyond fishes, partly because lenses in many non-fish taxa are softer and more hydrated, which complicates fine, repeatable sectioning. A pretreatment protocol that dries lenses before dividing them into numerous thin sections has been proposed, enabling the reconstruction of isotope ratio trajectories across the entire lifespan (Matsubayashi et al. 2025). This drying-based workflow has expanded sequential lens isotope analysis from fishes to other vertebrate groups, including birds (Hasegawa et al. 2025), mammals (Miura et al. 2025), and amphibians (Shiga et al. 2025). Moreover, the method enables each section to be aligned to the growth stage with higher precision; thus, it has opened new applications, such as reconstructing lifetime migration histories in fishes (Matsubayashi et al. 2025). Therefore, the use of eye-lens isotope chronologies is maximized when an appropriate and well-resolved temporal axis can be assigned to the sequence of lens sections.

Prolonged somatic growth nearly enables lifetime tracking of isotope histories in fishes and amphibians (Shiga et al. 2024; Matsubayashi et al. 2025). Skeletal growth markedly slows around sexual maturity in mammals (Castanet et al. 2004; Cubo et al. 2002); thus, eye-lens growth likely reaches an early asymptote, narrowing the recoverable time span. To support this interpretation, Miura et al. (2025) demonstrated that lens isotope ratios in brown bears capture lactation-to-weaning dietary shifts, with the highest temporal resolution largely confined to ∼1–2 years after birth and declining as lens growth slows. Hasegawa et al. (2025) demonstrated that lenses in birds retain pre-hatching isotope signatures and track early post-hatching dietary shifts; however, the lens size plateaus by ∼50 days; thus, the extent of later-life archiving remains unclear. Therefore, applying sequential eye lens isotope analysis to mammals and birds requires taxon-specific calibration of lens accretion timing, especially when growth decelerates, thereby limiting chronological resolution.

Radiocarbon (¹⁴C) chronometry provides a particularly powerful tool for constraining the effective time axis archived in eye lenses. In terrestrial systems, atmospheric nuclear weapons testing during the 1950s and the early 1960s caused a rapid and well-characterized increase in atmospheric ¹⁴C (the bomb pulse), which peaked around 1963 (Kutschera 2022). This bomb-pulse signal can function as a natural time marker for dating the carbon incorporated into animal eye lenses. However, applying bomb-pulse ¹⁴C to avian and mammalian lenses becomes complicated when aquatic-derived carbon contributes to the diet; freshwater food webs may incorporate ¹⁴C-depleted carbon (Rech et al. 2023), whereas marine food webs can demonstrate substantial phase lags associated with the atmospheric bomb pulse due to reservoir effects (Alves et al. 2018). Consequently, suitable target individuals should be demonstrably terrestrial and sufficiently old (on the order of decades) so that lens material spans a period in which the bomb-pulse variation is clearly represented. Obtaining such specimens from wild populations is generally challenging; however, individuals with well-documented captive husbandry histories can provide comparatively accessible and well-controlled samples for establishing lens-specific chronologies.

In this study, we measured sequential radiocarbon on eye-lens sections from multiple avian and mammalian species that died in zoological collections, intending to identify the chronological axis represented by isotope ratios archived in the lens. Most of the individuals analyzed in this study have lived for more than 10 years and have been maintained on diets free or largely free of aquatic-derived foods (Supplemental Text), thereby minimizing potential confounding from freshwater or marine carbon sources. By comparing radiocarbon profiles among species and across lens regions, we assessed how lens accretion rates and the duration of isotopic information retention differ among taxa and evaluated the validity and practical limits of using eye lenses for retrospective life-history reconstruction.

## 2 Methods/Experimental

### 2.1 Sample preparation

No animals were specifically sacrificed for this study. All animals analyzed were individuals that had died between 2011 and 2025 at Asa Zoo, Kyoto City Zoo, Chiba City Zoo, and Hirakawa Zoo in Japan (see Supplemental Text for details). Table 1 summarizes the age at death. Each facility permitted sample collection, and their respective institutional ethics committees approved all procedures (Approval numbers: ChbR6 [Chiba City Zoo], 2023-KCZ-004, 2024-KCZ-014 [Kyoto City Zoo]). The eyeballs were extracted and frozen at −30℃ at each zoo. Lens isolation was conducted according to established methods (Miura et al. 2025; Hasegawa et al. 2025), with slight modifications for certain species as described below.

**Table 1.** Comparison of calibrated 14C age and actual bi1th and death Year of zoo-kept animals.

| Common Name<br>/ Species Name | Year of birth or<br>immigration year<br>estimate | Years of Death | Age at Death | Calculated <sup>14</sup>C<br>Age of birth by<br>models | <sup>14</sup>C Age of Lens<br>Center (Mean ± SD) | <sup>14</sup>C Age of Lens<br>Outer (Mean ± SD) | Oldest Traceable<br>Record (Years) | Period Preserved in<br>Eye Lens (Year) |
| --- | --- | --- | --- | --- | --- | --- | --- | --- |
| Great green macaw<br>/ *Ara ambiguus* | AD 2011.6<br>(Birth) | AD 2011.9 | 0.3 | NA | cal AD 2011.7 ± 1.6 | cal AD 2012.1 ± 1.4 | 0.2 | 0.3 |
| Scarlet Macaw<br>/ *Ara macao* | AD 1997<br>(Birth) | AD 2024 | 27 | NA | cal AD 1997.6 ± 1.8 | cal AD 2002.8 ± 1.5 | 26.4 | 5.2 |
| Triton Cockatoo<br>/ *Cacatua galerita triton* | AD 2020<br>(Birth) | AD 2023 | 3 | NA | cal AD 2018.1 ± 1.7 | cal AD 2017.6 ± 2.1 | 5.0 | -0.5 |
| Triton Cockatoo<br>/ *Cacatua galerita triton* | AD 1977<br>(Immigration) | AD 2021 | 44 | 1977.7 | cal AD 1978.0 ± 1.0 | cal AD 1986.8 ± 1.2 | 43.0 | 9.1 |
| Sulphur-Crested Cockatoo<br>/ *Cacatua galerita* | AD 1975<br>(Immigration) | AD 2025 | 50 < | 1965.4 | cal AD 1968.7 ± 1.3 | cal AD 1996.1 ± 1.4 | 56.3 | 27.4 |
| Grey Peacockpheasant /<br>*Polyplectron bicalcaratum* | AD 2009<br>(Birth) | AD 2024 | 15 | 2009.3 | cal AD 2009.3 ± 2.1 | cal AD 2014.5 ± 1.9 | 14.7 | 5.2 |
| Mallard<br>/ *Anas platyrhynchos* | AD 2011<br>(Birth) | AD 2023 | 12 | 2011.2 | cal AD 2011.8 ± 1.5 | cal AD 2016.1 ± 2.4 | 11.2 | 4.3 |
| Ring-tailed lemur<br>/ *Lemur catta* | AD 1999<br>(Birth) | AD 2024 | 25 | 1999.9 | cal AD 2000.2 ± 1.6 | cal AD 2019.2 ± 0.0 | 23.8 | 19.1 |
| Siberian tiger / *Panthera*<br>*tigris altaica* | AD 2005<br>(Birth) | AD 2024 | 19 | 2002.3 | cal AD 2002.4 ± 1.6 | cal AD 2017.8 ± 2.0 | 21.6 | 15.4 |

In the Siberian tiger, tissue ranging from 0.24 to 3.39 mm from the lens nucleus was fragmented as a cohesive mass during isolation. Therefore, the central portion of the lens, where the location could be reliably identified (0–0.24 mm), was analyzed together with the outer separable portions. Similarly, in the greater cockatoo (bird 5), the central portion of the lens (0–1.697 mm from the center) was broken during the separation process; thus, the corresponding central region of the lens from the contralateral eye (0–1.43 mm) was used for analysis.

### 2.2 Radiocarbon Dating

Lens sections were subjected to graphitization at the Research Institute for Humanity and Nature and subsequently measured for radiocarbon (¹⁴C) concentration using an accelerator mass spectrometer at the Institute of Accelerator Analysis Ltd. (Kanagawa, Japan). The ^14^C/^12^C ratios were normalized using five replicates of National Institute of Standards and Technology (NIST) oxalic acid standards. Three replicates of chemical blanks were analyzed to assess the instrumental background. The grouped measurements of the NIST standard and chemical blanks were used as standard and background values, respectively. Background correction was applied to the measured ^14^C/^12^C ratios of the samples and standards. IAEA-C6 (ANU sucrose) was analyzed as a secondary standard for quality control. The measured F^14^C values are consistent with the certified value of IAEA-C6 (1.5061 ± 0.0011 F^14^C) within the combined uncertainty (2σ) of the measured and certified values. The normalized activity of the modern standard (A_ON_) and the sample activity (A_SN_) were calculated from the measured ^14^C/^12^C ratios (Mann 1983; Stuiver and Polach 1977). Radiocarbon concentrations are reported as fractional modern carbon (F^14^C), representing the relative concentration of radiocarbon compared with the atmospheric level before nuclear testing in AD 1950, where modern carbon is defined as F^14^C = 1.0 (Reimer et al. 2004). Fraction modern carbon (F^14^C) was calculated according to the definition of Jull et al. (2013) as follows:

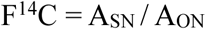

The CALIBomb program was used to estimate the calendar year of carbon fixation from the measured F^14^C values using the NHZ2 and NHZ3 (IntCal20) calibration datasets for the Northern Hemisphere summer tree-ring data and the SHZ3 (SHCal20) calibration datasets for the Southern Hemisphere data to account for variations in atmospheric ^14^C concentrations (Reimer et al. 2013; Reimer and Reimer 2025).

Calendar years corresponding to each lens section were subsequently identified (Table S1). Figure 1 was generated using the SHZ3 calibration curve, representing wide ranges in Indonesia, New Guinea, and Australia for birds 4 and 5, whose original habitats before introduction to Japan were unknown. Table S2 provides the calibration results using NHZ2 and NHZ3, corresponding to Japan and northern Indonesia. Differences in zone selection had minimal impact on age estimates, with a maximum deviation of ± 0.3 years (Table S2).

**Figure 1.**
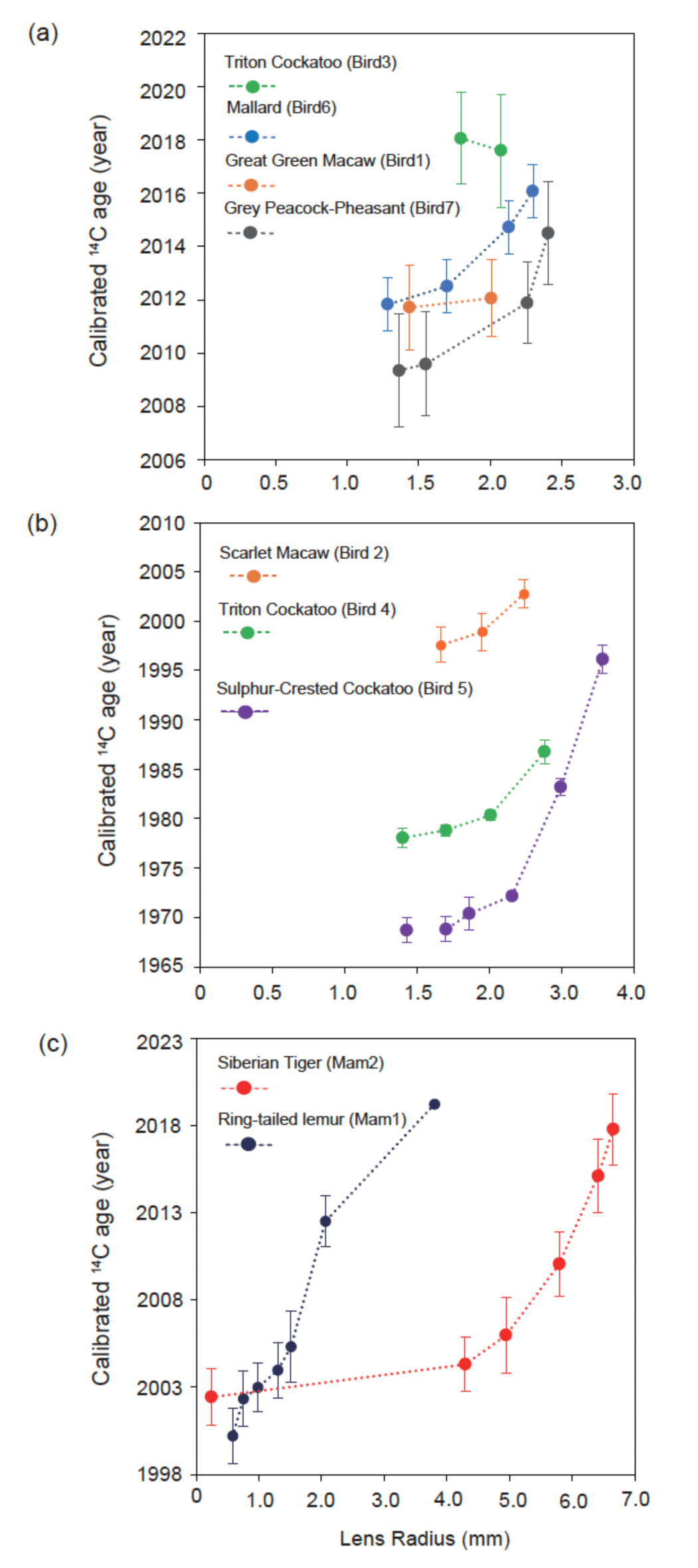
Calibrated ^14^C ages of eye lens sections from zoo-kept animals: (a) birds born after 2009, (b) birds born after 1997, and (c) mammals.

### 2.3 Lens Growth Model

Lens growth has been described in terms of weight, following either a monophasic or biphasic growth pattern (Augusteyn 2007, 2008, 2014a, 2014b; Bassnett and Sikic 2017). Based on an analysis of 130 lenses, Augusteyn (2014a) concluded that, except for six species, all lenses demonstrate monophasic growth in terms of weight. In this study, we assumed an exponential model based on lens weight to describe the association between age (identified by ¹⁴C dating) and position within the lens as follows:

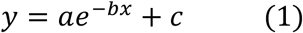

where *x* indicates the lens radius, *y* denotes the ¹⁴C-derived age, *a* represents a scaling coefficient, *b* demonstrates the decay rate, and *c* represents the initial value of the model. These parameters were estimated using the least-squares method. The Great Green Macaw (*Ara ambigua*, bird 1) and Triton Cockatoo (*Cacatua galerita triton*, bird 3) were excluded from this model estimation because bird 1 was only 4 months old and bird 3 lived for just 3 years, leading to insufficient data points for analysis. Lens growth for primates may follow a biphasic pattern (Augusteyn 2007), comprising an exponential growth model representing rapid growth during the prenatal period and the first few years postnatal, followed by a linear regression model (Mohamed and Augusteyn 2018). However, the data for the ring-tailed lemur demonstrated a growth pattern different from that of humans. Therefore, data were categorized into three growth phases according to the visual inspection of the scatterplot: an exponential model representing rapid early growth (2000.2–2002.3 year) (phase 1), a slower exponential growth model than the initial phase (2002.3–2012.5 year) (phase 2), and a more gradual linear regression model (2012.5–2019.3 year) (phase 3). In phase 1, the exponential model of equation (1) was fitted by selecting the initial value C within the range from the birth year (1999 year) to the oldest ¹⁴C-estimated age in this phase (2000.2 year) to find the model that best fit the data using the least squares method (the model with the minimum residual sum of squares was selected). In phase 2, the exponential model of equation (1) was fitted by selecting C within the range from 1999 to the oldest ¹⁴C-estimated age in this phase (2002.3 year), and the best-fitting model was identified using the least-squares method. In phase 3, a linear regression between the ¹⁴C ages and lens positions (radius) was fitted. Conducting statistical tests to rigorously evaluate the fit of the models to the data was impossible due to the limited sample size. The residual sum of squares and coefficient of determination (R²) were used as quantitative indicators of model fit and calculated using R (R Core Team 2024).

## 3 Results

### 3.1 ^14^C age of the lens nucleus

In both mammals and birds, the calibrated ¹⁴C ages increased from the eye lens center toward the periphery (Figure 1). The central part of the eye lens demonstrated the oldest ¹⁴C age, with ages approaching the present toward the outer regions. Sections from the central portion of the eye lens yielded ¹⁴C ages that closely matched the known birth years (Table 1). The ¹⁴C age in the mammalian species was AD 2002.4 ± 1.6 year for a Siberian tiger born in 2005 and AD 2000.2 ± 1.6 year for a ring-tailed lemur (*Lemur catta*) born in 1999. In the avian species, the scarlet macaw (*Ara macao*) born in 1997 exhibited a ¹⁴C age of AD 1997.6 ± 1.8 years, corresponding to its birth year. For more recent samples, the ¹⁴C age of a mallard (*Anas platyrhynchos*) born in 2011 was AD 2011.8 ± 1.5 years, and a gray peacock-pheasant (*Polyplectron bicalcaratum*) born in 2009 was AD 2009.3 year, closely matching the actual birth year. ¹⁴C ages for birds with unknown import years, including a sulphur-crested cockatoo (*Cacatua galerita*) imported before 1975 and a blue-eyed cockatoo imported in 1977, were AD 1968.7 ± 1.3 years and AD 1978.0 ± 1.0 years in the central eye lens, respectively, consistent with being born in or before the import year. Conversely, no ages more recent than 2018 were detected for triton cockatoos (*Cacatua galerita triton*) born after 2020 (2020–2023) (AD 2018.1 ± 1.7 and AD 2017.6 ± 2.1 years), leading to a discrepancy with the actual birth years.

### 3.2 Lens Growth Model

The association between ¹⁴C age and radial position in the eye lens was well fitted by an exponential model in both taxonomic groups (R^2^ > 0.99, Figure 2(a)–(e)). Using the exponential models, the calculated birth years (i.e., the initial values of the model, c; see Methods) were calculated and closely matched the known age of birth: AD 2002.3 for the tiger, AD 2009.3 for the gray peacock-pheasant, and AD 2011.2 for the mallard (Table 1). The calculated birth years for imported birds with unknown birth records were AD 1977.7 for the blue-eyed cockatoo and AD 1965.4 for the sulphur-crested cockatoo.

**Figure 2.**
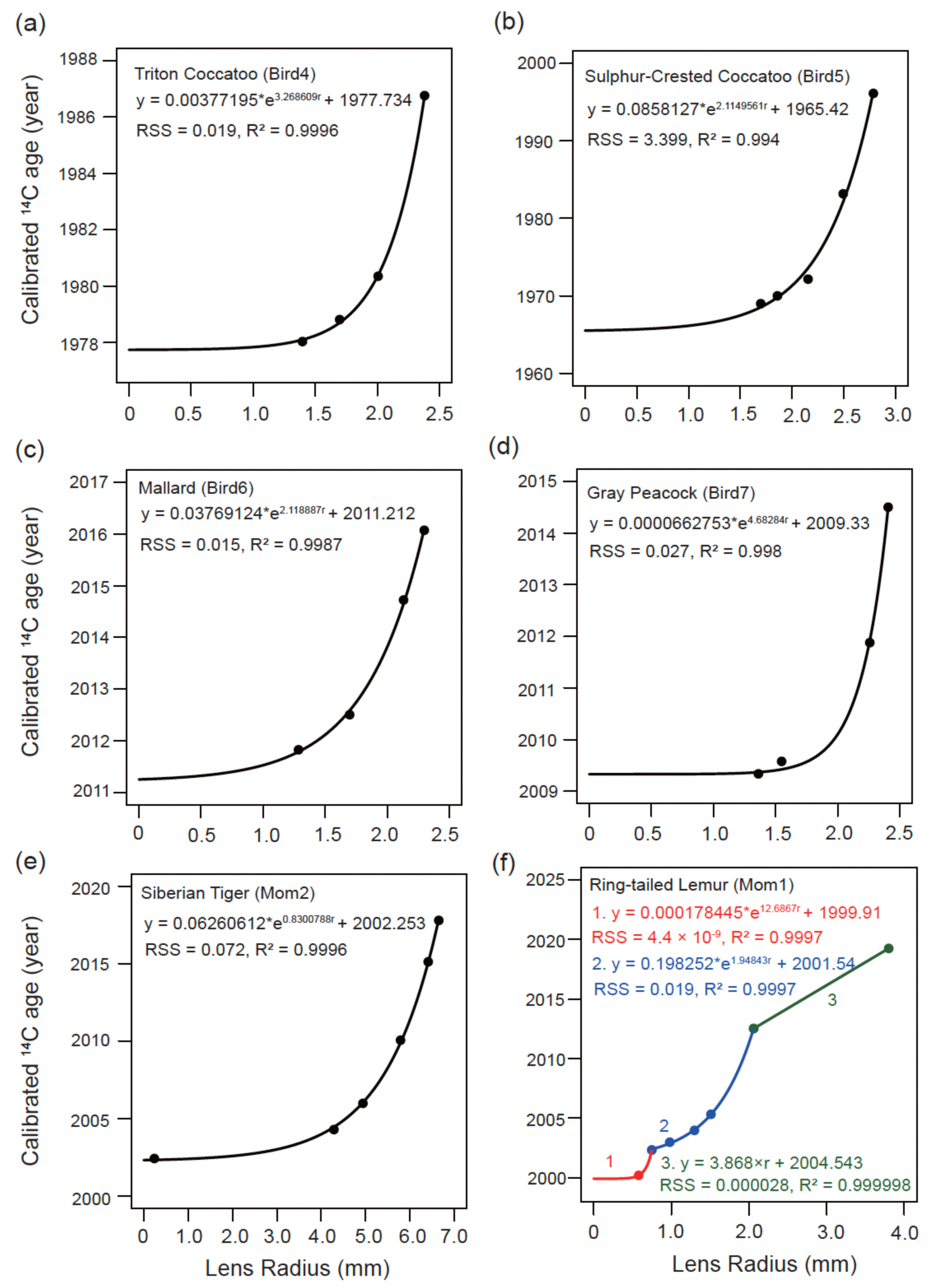
Exponential model ((a)-(e)) and multiple model (f) demonstrating the association between the calibrated ^14^C age (y) and eye lens radius (r) in zoo-kept animals.

The ring-tailed lemur demonstrated a distinct lens growth pattern described by a three-phase model: between AD 2000.2 and AD 2002.3, between AD 2002.3 and AD 2012.5, and between AD 2012.5 and AD 2019.2 (Figure 2 (f)). The calculated birth year (AD 1999.9) closely matched its actual birth (1999) (Figure 2 (f)).

### 3.3 Temporal window recorded in the eye lens

In mammals, the 25-year-old ring-tailed lemur enabled reconstruction over 19.1 years, and the 19-year-old Amur tiger over 15.4 years (Table 1). Both animals died in 2024 and were reconstructed up to AD 2019.2 for the lemur and AD 2017.8 for the tiger in the outermost layer. In the avian species evaluated, the reconstructed time span considerably varied among individuals and was generally shorter than in the two-mammalian species (Table 1). The calibrated ^14^C age for the scarlet macaw (1997–2024) in the central eye lens was AD 1997.6 ± 1.6 years and the outermost layer AD 2002.8 ± 1.5 years, leading to a 5-year reconstructable period. The reconstructable range for the blue-eyed cockatoo (imported to Japan in 1977, died 2021) was AD 1978.0 to 1986.8, spanning 9 years. The longest reconstruction was achieved in the sulphur-crested cockatoo (donated to the zoo in 1975, died 2025), covering AD 1968.7 to AD 1996.1, spanning 27.4 years. Only 4.3 years could be reconstructed for more recently born individuals, such as the mallard (2011–2023) (up to 2016), whereas only 5.2 years for the gray peacock-pheasant (2009–2024) (up to 2014.5), similar to mammals, indicating that ages close to the present could not be resolved using the radiocarbon calibration approach applied in this study.

## 4 Discussion

### 4.1 The validity of time-series isotope reconstruction in mammal and bird eye lenses

In both mammals and birds, ¹⁴C ages increased from the eye lens center toward the periphery (Figure 1), indicating that the eye lens preserves the chronological order of formation, with older cells at the central part of the lens (i.e., lens nucleus) and newer cells toward the outer regions. Such a temporal gradient from the center toward the periphery is also consistent with the structural characteristics of the lens (Priolo et al. 1999). Cells along the optical pathway must remain free of light-scattering structures, such as organelles, throughout life, to maintain lens transparency. During lens development, cell fusion leading to the formation of a syncytial structure has been reported in the lens core but not in the peripheral and outer regions of the lens (Shestopalov and Bassnett 2000). Our results support the hypothesis that the syncytial organization is restricted to the central region of the lens and further indicate that the lens comprises a centrally formed core established before birth and surrounded by outer membrane structures that do not undergo cell fusion but instead retain newly accumulated information sequentially over time.

The ^14^C dates obtained from the central region of the lens were consistent with the known birth year, providing further support for the robustness of our method. This study provides the first retrospective reconstruction of birth dates from approximately 20 years ago in non-human mammals and 50 years ago in birds. Age estimation in birds has traditionally relied on mark–recapture records obtained from individuals marked at birth, making it extremely challenging to identify the ages of long-lived individuals. Our results demonstrate that the lens provides a novel and reliable archive for age estimation in birds, providing a new approach to identifying the ages of long-lived individuals.

### 4.2 Temporal Model of the Eye Lens

Most of the age–radius associations along the eye lens growing axis (Figure 2) were consistent with previously reported weight-based growth curves (Augusteyn 2007, 2008, 2014a, 2014b; Bassnett and Sikic 2017) and expressed as exponential models (Figure 2 (a)-(e)), except for the ring-tailed lemur (Figure 2 (f)). Previous studies have shown that primate lens growth varies from that of other mammals (Augusteyn 2014a) and is likely expressed as a two-phase model (Augusteyn 2007; Mohamed & Augusteyn 2018). However, our analyses revealed that a three-phase model fit the lens growth pattern better than the conventional two-phase model (Figure 2f). This resembles body-mass–based growth trajectories reported for many Lemuridae, partitioned into an initial period of rapid growth, a subsequent and clearly distinguishable period of slower growth, and a final phase in which growth nearly ceases (Tennenhouse 2015). Accordingly, lens growth dynamics may share greater similarities with body-mass–based growth models than with skeletal-length–based growth models. Therefore, a universally exponential lens growth pattern should not be assumed when interpreting the time axis of serial eye-lens isotope ratios. Instead, the lens-based chronology should be interpreted and parameterized in a species-specific manner, informed by ontogenetic shifts in body mass and related growth dynamics.

### 4.3. Limitations in temporal resolution at the outer region of the eye lens

Time-series isotope reconstruction using the ocular lens indicated that temporal resolution decreases toward the outer layers of the lens. The exponential age–radius associations obtained in this study (Figure 2) indicate that the amount of newly accumulated protein per unit time rapidly decreases toward the lens periphery, and this structural characteristic likely imposes a fundamental limitation on chronological reconstruction in the outer regions. Particularly, peripheral lens sections may contain proteins formed over multiple years, causing isotopic values to be averaged across that interval and thereby reducing temporal resolution.

Despite this decline in resolution in mammals, records were retained until relatively recent years. Approximately 15 years of reconstruction were achieved in the tiger and 19 years in the ring-tailed lemur, with the calibrated ¹⁴C ages of the outermost layers reaching close to the analytical limit around 2018. Therefore, year-scale temporal averaging may occur in the outer lens layers; however, the capacity for long-term information retention reported in humans may be present in non-human mammals. The failure of the outermost layers to reach the year of death may reflect the averaging effect related to lens sectioning and/or the analytical limitation of CALIBomb near the present. Therefore, the present data hindered us from fully distinguishing limitations in biological recording from those in radiocarbon age assignment. Analyses of individuals that died before the analytical limit will be necessary to further assess the true reconstruction limits of mammalian lenses.

Conversely, birds demonstrated not only reduced temporal resolution in the outer layers but also clear limitations in the total reconstructable period itself. The calibrated ¹⁴C ages in the outermost layer only reached 2003 in the Scarlet Macaw (1997–2024), whereas reconstruction extended only to 1987 in the Triton Cockatoo (1977–2021). Reconstruction only reached 1996 even in the Sulphur-crested Cockatoo, which demonstrated the longest reconstruction period (introduced in 1975 and died in 2025). These results indicate that, in birds, information immediately preceding death is not retained and that the amount of newly recorded information in the outer lens layers is inherently limited.

This interpretation is consistent with previous studies demonstrating that lens weight in the House Sparrow reaches a plateau approximately 50 days after hatching, after which age estimation using the lens becomes challenging. Lens growth in birds may cease or slow dramatically relatively early in life, thereby limiting the accumulation of new proteins and reducing the ability of the lens to preserve long-term chronological records.

## 5 Conclusions

In this study, radiocarbon dating of eye-lens sections from mammals and birds was utilized to reconstruct the temporal framework preserved within the eye lens. Our results in the mammalian species examined in this study indicate that the eye lens may retain long-term isotopic information over much of the lifespan, extending beyond previously assumed temporal limits. Continuous lifetime records may not be fully preserved in birds; however, isotopic information from birth and early life stages was retained over extended periods.

Overall, our results indicate that the temporal structure recorded in eye lenses is shaped by species-specific growth dynamics. The onset and rate of lens growth attenuation strongly influenced both temporal resolution and duration of recoverable records. Expanding this approach to a wider range of taxa with diverse life-history strategies is crucial for developing a more comprehensive understanding of how temporal information is archived in biological tissues.

## Declarations

### Availability of data and material

All data are available in the main text and the supplementary materials.

## Authors’ contributions

JM proposed the topic the study. EH conceived and designed the study. KM, YY, EH carried out the dissection of eye lenses. EH carried out the CO_2_ purification and Graphitization of samples, and JM and IT helped this experiment. EH analyzed the data and JM, KM, IT helped in their interpretation. JM and KM collaborated with the corresponding author in the construction of manuscript. All authors read and approved the final manuscript.

## Authors’ information

EH was employed by Research Institute for Humanity and Nature as a Restart Postdoctoral Fellowship (RPD) researcher.

## Acknowledgements

We are grateful for Dr. Wataru Anzai, Asa zoological Park, Kyoto city zoo, Chiba zoological park, Hirakawa zoological Park, and Maruyama zoological Park for the contributions for sample collection. We thank Research Institute for Humanity and Nature for Utilization of experimental facilities. We thank the Grant-in-Aid for JSPS Research Fellow JP23K2160.

