## Supplementary Text, Tables S1 to S2 for "Radiocarbon dating of bird and mammal eye lenses reveals long-term chronology"

**The PDF file includes:**

Supplementary Text
Tables S1 to S2

**Supplementary Text**

Animal History and Diet Information

**Bird 1**

*Species*: Great green macaw (*Ara ambigua*)
*History:* Born in July 2011 at Asa Zoological Park (Hiroshima, Japan).
Passed away at Asa Zoological Park in November 2011.
*Diet:* Terrestrial plant-based food.
*Zoning for CALIBomb calibration*: NHZ2.

**Bird 2**

*Species:* Scarlet macaw (*Ara macao*)
*History:* Born in 1997 at Asa Zoological Park.
Passed away at Asa Zoological Park in 2024.
*Diet:* Terrestrial plant-based food.
*Zoning for CALIBomb calibration*: NHZ2.

**Bird 3**

*Species:* Triton Cockatoo (*Cacatua galerita triton*).
*History:* Born in 2020 at Asa Zoological Park.
Passed away at Asa Zoological Park in 2023.
*Diet:* Terrestrial plant-based food.
Mazuri Large Bird Diet
https://mazuri.com/products/mazuri-large-bird-diet
Mazuri Parrot Breeder
https://mazuri.com/products/mazuri-parrot-brdr-diet?icid=recs_pdp_2
Grains and seeds: sunflower seeds, hemp seeds, corn, sorghum, pumpkin seeds, peanuts, white sunflower seeds
Fruits: apple, banana, mandarin orange, or orange
Root vegetables: sweet potato, carrot
Bread: white bread
*Zoning for CALIBomb calibration*: NHZ2.

**Bird 4**

*Species:* Triton Cockatoo (*Cacatua galerita triton*)
*Home Range:* New Guinea, Australia
*History:* Introduced to Japan from Indonesia in 1977. No information before 1977.
Donated to Asa Zoological Park in 1993.
Passed away at Asa Zoological Park in 2021.
*Diet:* Terrestrial plant-based food.
Mazuri Large Bird Diet
https://mazuri.com/products/mazuri-large-bird-diet
Mazuri Parrot Breeder
https://mazuri.com/products/mazuri-parrot-brdr-diet?icid=recs_pdp_2
Grains and seeds: sunflower seeds, hemp seeds, corn, sorghum, pumpkin seeds, peanuts, white sunflower seeds

Fruits: apple, banana, mandarin orange, or orange
Root vegetables: sweet potato, carrot
Bread: white bread

*Zoning for CALIBomb calibration*: Center of the lens. We used NHZ2, NHZ3, and SHZ3 as shown in Table S2. From the year bird4 came to Japan onwards, it is classified as NHZ2.

**Bird 5**
*Species:* Sulphur-crested Cockatoo (*Cacatua galerita*)
*History:* The exact date of arrival in Japan is unknown. A bird kept as a pet by a Kyoto resident was donated to the Kyoto City Zoo in 1974. Passed away in Kyoto City in 2025.
*Diet:* Terrestrial plant-based food.
Mazuri Large Bird Diet
https://mazuri.com/products/mazuri-large-bird-diet
Mazuri Parrot Breeder
https://mazuri.com/products/mazuri-parrot-brdr-diet?icid=recs_pdp_2
Grains and seeds: sunflower seeds, hemp seeds, corn, sorghum, pumpkin seeds, peanuts, white sunflower seeds
Fruits: apple, banana, mandarin orange, or orange
Root vegetables: sweet potato, carrot
Bread: white bread
*Zoning for CALIBomb calibration*: Center of the lens. We used NHZ2, NHZ3, and SHZ3 as shown in Table S2. From the year bird4 came to Japan onwards, it is classified as NHZ2.

**Bird 6**
*Species:* Mallard (*Anas platyrhynchos*)
*History:* Born in 2011 at Chiba City Zoological Park.
Passed away at Chiba City Zoological Park in 2023.
*Diet:* Terrestrial-derived food.
*Zoning for CALIBomb calibration*: NHZ2.

**Bird 7**
*Species:* Grey peacock pheasant (*Polyplectron bicalcaratum*)
*History:* Born in 2009 at Chiba City Zoo. Passed away at Chiba City Zoo in 2024.
*Diet:* Terrestrial-derived food.
*Zoning for CALIBomb calibration*: NHZ2.

**Mam 1**
*Species:* Ring-tailed Lemur *Lemur catta*
*History:* Born in 1999 in Hyogo Prefecture.
Passed away at Kyoto City Zoo in 2024.

*Diet:* Mainly terrestrial-derived food.Feed composition: Oriental Yeast Old World Monkey pellets (10 kg) (Ingredients: wheat flour, corn, soybean meal, skimmed milk powder, fish meal, peanut meal, vitamins, minerals).
*Zoning for CALIBomb calibration*: NHZ2.

**Mam2**
*Species:* Siberian Tiger *Panthera tigris altaica*
*History:* Born in 2005 in Shizuoka Prefecture.
Passed away in Kyoto City in 2024.
*Diet:* Terrestrial-derived food. Carnivorous.
*Zoning for CALIBomb calibration*: NHZ2.

**Table S1. $F^{14}C$ values of each eye lens section at different distances from the lens center (Radius).**

| ID | Species Name | Radius (mm) | $F^{14}C$ (Mean±SD) |
|---|---|---|---|
| Bird1 | *Ara ambiguus* | 2.0 | 1.0386±0.0028 |
| Bird1 | *Ara ambiguus* | 1.4 | 1.0410±0.0028 |
| Bird2 | *Ara macao* | 2.2 | 1.0800±0.0028 |
| Bird2 | *Ara macao* | 1.9 | 1.1002±0.0028 |
| Bird2 | *Ara macao* | 1.7 | 1.1074±0.0029 |
| Bird3 | *Cacatua galerita triton* | 2.1 | 1.0152±0.0027 |
| Bird3 | *Cacatua galerita triton* | 1.8 | 1.0127±0.0027 |
| Bird4 | *Cacatua galerita triton* | 2.4 | 1.1917±0.003 |
| Bird4 | *Cacatua galerita triton* | 2.0 | 1.2761±0.0031 |
| Bird4 | *Cacatua galerita triton* | 1.7 | 1.3188±0.0032 |
| Bird4 | *Cacatua galerita triton* | 1.4 | 1.3291±0.0032 |
| Bird5 | *Cacatua galerita* | 2.8 | 1.115±0.0028 |
| Bird5 | *Cacatua galerita* | 2.5 | 1.2345±0.003 |
| Bird5 | *Cacatua galerita* | 2.2 | 1.4967±0.0034 |
| Bird5 | *Cacatua galerita* | 1.9 | 1.54±0.0035 |
| Bird5 | *Cacatua galerita* | 1.7 | 1.5682±0.0035 |
| Bird5 | *Cacatua galerita* | 1.4 | 1.5705±0.0035 |
| Bird6 | *Anas platyrhynchos* | 2.3 | 1.0212±0.0027 |
| Bird6 | *Anas platyrhynchos* | 2.1 | 1.0259±0.0027 |
| Bird6 | *Anas platyrhynchos* | 1.7 | 1.0359±0.0027 |
| Bird6 | *Anas platyrhynchos* | 1.3 | 1.0403±0.0028 |
| Bird7 | *Polyplectron bicalcaratum* | 2.4 | 1.0273±0.0027 |
| Bird7 | *Polyplectron bicalcaratum* | 2.3 | 1.0406±0.0028 |
| Bird7 | *Polyplectron bicalcaratum* | 1.5 | 1.0502±0.0027 |
| Bird7 | *Polyplectron bicalcaratum* | 1.4 | 1.0512±0.0028 |
| Mam1 | *Lemur catta* | 3.8 | 1.0013±0.0027 |
| Mam1 | *Lemur catta* | 2.1 | 1.0358±0.0026 |
| Mam1 | *Lemur catta* | 1.5 | 1.0676±0.0028 |
| Mam1 | *Lemur catta* | 1.3 | 1.0734±0.0028 |
| Mam1 | *Lemur catta* | 1.0 | 1.0781±0.0029 |
| Mam1 | *Lemur catta* | 0.8 | 1.0819±0.0028 |
| Mam1 | *Lemur catta* | 0.6 | 1.0932±0.0028 |
| Mam2 | *Panthera tigris altaica* | 6.7 | 1.0148±0.0027 |
| Mam2 | *Panthera tigris altaica* | 6.4 | 1.0241±0.0027 |
| Mam2 | *Panthera tigris altaica* | 5.8 | 1.0479±0.0027 |
| Mam2 | *Panthera tigris altaica* | 4.9 | 1.0634±0.0028 |
| Mam2 | *Panthera tigris altaica* | 4.3 | 1.0715±0.0028 |
| Mam2 | *Panthera tigris altaica* | 0.2 | 1.0815±0.0029 |

### Table S2. Birth and dead site and zone for $^{14}C$ Age

| ID | Species Name | Years of Birth or immigration | Birth site | Year of Death | Dead site | Zone | $^{14}C$ Age (Mean ±SD) | Lens Radius (mm) |
|---|---|---|---|---|---|---|---|---|
| Bird1 | *Ara ambiguus* | 2011 | Hiroshima | 2011 | Hiroshima | NHZ2 | 2012.1 ±1.4 | 2.0 |
| Bird1 | *Ara ambiguus* | 2011 | Hiroshima | 2011 | Hiroshima | NHZ2 | 2011.7±1.6 | 1.4 |
| Bird2 | *Ara macao* | 1997 | Hiroshima | 2024 | Hiroshima | NHZ2 | 2002.8±1.5 | 2.2 |
| Bird2 | *Ara macao* | 1997 | Hiroshima | 2024 | Hiroshima | NHZ2 | 1998.9±1.9 | 1.9 |
| Bird2 | *Ara macao* | 1997 | Hiroshima | 2024 | Hiroshima | NHZ2 | 1997.6±1.8 | 1.7 |
| Bird3 | *Cacatua galerita triton* | 2020 | Hiroshima | 2023 | Hiroshima | NHZ2 | 2017.6±2.1 | 2.1 |
| Bird3 | *Cacatua galerita triton* | 2020 | Hiroshima | 2023 | Hiroshima | NHZ2 | 2018.1±1.7 | 1.8 |
| Bird4 | *Cacatua galerita triton* | 1977 | NA | 2021 | Hiroshima | NHZ2 | 1986.8±1.2 | 2.4 |
| Bird4 | *Cacatua galerita triton* | 1977 | NA | 2021 | Hiroshima | NHZ2 | 1980.3±0.5 | 2.0 |
| Bird4 | *Cacatua galerita triton* | 1977 | NA | 2021 | Hiroshima | NHZ2<br>/NHZ3<br>/SHZ3 | 1978.8±0.6<br>1978.8±0.6<br>1978.7±0.8 | 1.7 |
| Bird4 | *Cacatua galerita triton* | 1977 | NA | 2021 | Hiroshima | NHZ2<br>/NHZ3<br>/SHZ3 | 1978.2±0.9<br>1978.2±0.9<br>1978.0±1.0 | 1.4 |
| Bird5 | *Cacatua galerita* | 1975 | NA | 2025 | Kyoto | NHZ2 | 1996.1±1.4 | 2.8 |
| Bird5 | *Cacatua galerita* | 1975 | NA | 2025 | Kyoto | NHZ2 | 1983.2±0.8 | 2.5 |
| Bird5 | *Cacatua galerita* | 1974 | NA | 2025 | Kyoto | NHZ2<br>/NHZ3<br>/SHZ3 | 1971.6±1.1<br>1972.2±0.3<br>1972.2±0.4 | 2.2 |
| Bird5 | *Cacatua galerita* | 1974 | NA | 2025 | Kyoto | NHZ2<br>/NHZ3<br>/SHZ3 | 1969.8±0.6<br>1970.4±1.7<br>1970.0±0.8 | 1.9 |
| Bird5 | *Cacatua galerita* | 1974 | NA | 2025 | Kyoto | NHZ2<br>/NHZ3<br>/SHZ | 1968.8±0.7<br>1968.8±1.3<br>1969.0±0.6 | 1.7 |
| Bird5 | *Cacatua galerita* | 1974 | NA | 2025 | Kyoto | NHZ2<br>/NHZ3<br>/SHZ | 1968.7±0.7<br>1968.7±1.3<br>1968.9±0.7 | 1.4 |
| Bird6 | *Anas platyrhynchos* | 2011 | Chiba | 2023 | Chiba | NHZ2 | 2016.1±2.4 | 2.3 |
| Bird6 | *Anas platyrhynchos* | 2011 | Chiba | 2023 | Chiba | NHZ2 | 2014.7±2.0 | 2.1 |
| Bird6 | *Anas platyrhynchos* | 2011 | Chiba | 2023 | Chiba | NHZ2 | 2012.5±1.5 | 1.7 |
| Bird6 | *Anas platyrhynchos* | 2011 | Chiba | 2023 | Chiba | NHZ2 | 2011.8±1.5 | 1.3 |
| Bird7 | *Polyplectron bicalcaratum* | 2009 | Chiba | 2024 | Chiba | NHZ2 | 2014.5±1.9 | 2.4 |
| Bird7 | *Polyplectron bicalcaratum* | 2009 | Chiba | 2024 | Chiba | NHZ2 | 2011.9±1.5 | 2.3 |
| Bird7 | *Polyplectron bicalcaratum* | 2009 | Chiba | 2024 | Chiba | NHZ2 | 2009.6±2.0 | 1.5 |
| Bird7 | *Polyplectron bicalcaratum* | 2009 | Chiba | 2024 | Chiba | NHZ2 | 2009.3±2.1 | 1.4 |
| Mam1 | *Lemur catta* | 1999 | Hyogo | 2024 | Kyoto | NHZ2 | 2019.2±0.0 | 3.8 |
| Mam1 | *Lemur catta* | 1999 | Hyogo | 2024 | Kyoto | NHZ2 | 2012.5±1.5 | 2.1 |
| Mam1 | *Lemur catta* | 1999 | Hyogo | 2024 | Kyoto | NHZ2 | 2005.3±2.0 | 1.5 |
| Mam1 | *Lemur catta* | 1999 | Hyogo | 2024 | Kyoto | NHZ2 | 2004.0±1.6 | 1.3 |
| Mam1 | *Lemur catta* | 1999 | Hyogo | 2024 | Kyoto | NHZ2 | 2003.0±1.4 | 1.0 |
| Mam1 | *Lemur catta* | 1999 | Hyogo | 2024 | Kyoto | NHZ2 | 2002.3±1.6 | 0.8 |
| Mam1 | *Lemur catta* | 1999 | Hyogo | 2024 | Kyoto | NHZ2 | 2000.2±1.6 | 0.6 |

| | | | | | | | | |
|---|---|---|---|---|---|---|---|---|
| Mam2 | *Panthera tigris altaica* | 2004 | Shizuoka | 2024 | Kyoto | NHZ2 | 2017.8±2.0 | 6.7 |
| Mam2 | *Panthera tigris altaica* | 2004 | Shizuoka | 2024 | Kyoto | NHZ2 | 2015.1±2.1 | 6.4 |
| Mam2 | *Panthera tigris altaica* | 2004 | Shizuoka | 2024 | Kyoto | NHZ2 | 2010.1±1.8 | 5.8 |
| Mam2 | *Panthera tigris altaica* | 2004 | Shizuoka | 2024 | Kyoto | NHZ2 | 2006.0±2.2 | 4.9 |
| Mam2 | *Panthera tigris altaica* | 2004 | Shizuoka | 2024 | Kyoto | NHZ2 | 2004.3±1.6 | 4.3 |
| Mam2 | *Panthera tigris altaica* | 2004 | Shizuoka | 2024 | Kyoto | NHZ2 | 2002.4±1.6 | 0.2 |
